# How drugs modulate the performance of the human heart

**DOI:** 10.1101/2021.07.12.452066

**Authors:** Mathias Peirlinck, Jiang Yao, Francisco Sahli Costabal, Ellen Kuhl

## Abstract

Many drugs interact with ion channels in the cells of the heart and trigger heart rhythm disorders with potentially fatal consequences. Computational modeling can provide mechanistic insight into the onset and propagation of drug-induced arrhythmias, but the effect of drugs on the mechanical behavior of the heart remains poorly understood. Here we establish a multiphysics framework that integrates the biochemical, electrical, and mechanical effects of drugs from single cardiac cells to the overall response of the whole heart. For the example of the drug dofetilide, we show that drug concentrations of 3.0x and 4.8x increase the heart rate to 122 and 114 beats per minute, increase the myofiber stretches up to 10%, and decrease tissue relaxation by 6%. Strikingly, the drug-induced interventricular and atrial-ventricular dyssynchrony results in a 2.5% decreased and 7% increased cardiac output, respectively. Our results demonstrate the potential for multiphysics, multiscale modeling towards understanding the mechanical implications of drug-induced arrhythmias. Knowing how differing drug concentrations affect the performance of the heart has important clinical implications in drug safety evaluation and personalized medicine.

## Motivation

All medications have side effects. Drug-induced ventricular arrhythmia and sudden cardiac death are rare but severe adverse events that should be avoided at all cost. Consequently, when a new drug is developed, the proarrhythmic potential of the new compounds is a key concern. The current gold standard pharmacological pro-arrhythmic risk stratification combines in vitro experiments to quantify pharmacological blocking of specific cardiac ion channels, with electrocardiographic large animal experiments and clinical studies focusing on changes in tissue activation duration. Although these biomarkers show good sensitivity, they are costly and have poor specificity, potentially blocking safe new drugs from ever reaching the market (38). To develop novel and more accurate drug-induced arrhythmia biomarkers, multiphysics multiscale models mechanistically couple what a pharmacologist sees in a single cell experiment to what a physician sees in a clinical electrocardiogram (7). As part of these efforts, our group has recently proposed an electrophysiological exposure-response simulator that integrates the interaction between multiple drug compounds and specific ionic currents at the cellular scale with the intrinsic cardiac anisotropic conductivity at the tissue scale and the transmural heterogeneity and tissue organization at the organ scale (42). This framework allows us to conduct in silico drug trials for multiple drugs at various concentrations (46), providing risk categories that correlate well with reported drug-induced arrhythmia incidence (43). Based on these results, we trained and validated a binary risk classifier that accurately predicts the critical pro-arrhythmic drug concentration (45). From a clinical perspective however, a binary risk classification only provides a limited insight into the malignancy of the arrhythmic event. Dependent on the periodicity of the drug-induced arrhythmia, the cardiac output can increase, decrease or stay relatively constant. Consequently, short-duration non-sustained arrhythmogenicity can have multiple outcomes for the patient. In this study, we extended our framework to provide insights into the changing cardiac output of the heart at varying arrhythmogenic drug concentrations. More specifically, we use the electrophysiogical activation sequence to drive biomechanical tissue contraction in the human heart and study the resulting hemodynamic effects on the whole-body cardiovascular circulation. Doing so, we compute a drug’s pharmacological potential to impede efficient propulsion of blood through the heart chambers and the rest of the body. As such, we extend what a physician sees in a clinical electrocardiogram to what a patient feels and how likely they are to survive specific dosage-dependent drug-induced arrhythmia events.

## Methods

### Cardiac electrophysiology

We simulate the electrophysiological behavior of cardiac tissue using the monodomain model (41). The main variable of the monodomain model is the transmembrane potential *ϕ*, the difference between the intra- and extra-cellular potentials. The transmembrane potential is governed by a reaction-diffusion equation (20)

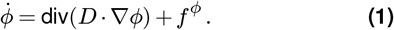

Here, we introduce the source term *f^ϕ^* which represents the ionic currents across the cell membrane and the conductivity tensor *D*, which we further decompose into fast *D*^||^ and slow *D*^⊥^ signal propagation parallel and perpendicular to the cardiac mucle fiber direction *f* respectively (12),

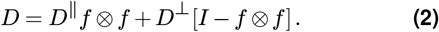

In general, the ionic currents *f^ϕ^*(*ϕ,q*(*ϕ*);*t*) are functions of the transmembrane potential *ϕ* and a set of state variables *q*(*ϕ*) (22; 53), where the state variables themselves are governed by ordinary differential equations, 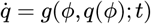. The number of currents and state variables determines the complexity of the model and varies for different cell types. To simulate the electrophysiological behavior of the Purkinje fiber network, we choose the Stewart model for human Purkinje fiber cells (47). A characteristic feature of this model is the automaticity of its action potential, which enables the cells to self-excite without an external stimulus. This model is based on 14 ionic currents

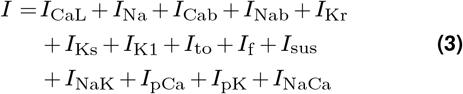

that are defined through 20 state variables. To study the spatiotemporal action potential evolution in the myocardium, we select the O’Hara-Rudy model for human ventricular cardiomyocytes (27). This model was developed based on a vast amount of human experimental data and includes description of key ionic currents for drug-induced arrhythmias. More specifically, the model is based on 15 ionic currents,

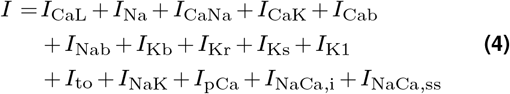

of which we replaced the fast sodium current *I*_Na_ of the original O’Hara-Rudy model with a modified fast sodium current of the ten Tusscher model (48) to model propagation in tissue scale simulations (36). These 15 transmembrane ion currents are defined through a total of 39 state variables. To account for regional specificity, we reparametrize the cardiomyocyte cell model for three different cell types: endocardial, mid-wall, and epicardial cells (27).

We incorporate drug effects by blocking the currents of the pharmacologically affected ion channels on the Purkinje and cardiomyocyte cell membrane. Based on discrete experimental patch clamp measurements of the fractional ion channel block at various drug concentrations (8), we fit a Hill-type equation

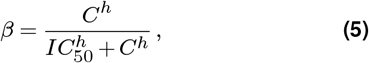

to describe fractional blockage *β* at any possible drug concentrations *C*. Here, the drug’s concentration-specific ion channel block is completely described by two parameters: the exponent *h* and the concentration *IC*_50_ required to achieve a 50% current block. We focus on the drug dofetilide, an anti-arrhythmic drug typically used for treating atrial fibrillation. This drug is a selective *I*_Kr_ blocker, characterized by the Hill parameters *h*_Kr_ = 0.65 and *IC*_50,Kr_ = 1.55 nM. To apply the drug at a desired concentration *C*, we calculate the fractional blockage *β*_Kr_ and scale the rapid delayed rectifier potassium ion channel conductance,

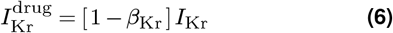

by multiplying the baseline current *I*_Kr_ with the fractional blockage [1 – *β*_Kr_]. Based on previous work which delineated the critical concentrations of dofetilide for developing arrhythmic events (44), we focus in particular on applying dofetilide at 3x, 4.8x, and 18.5x its free plasma concentration, *C*^max^ = 2.1 nM. This corresponds to dofetilide concentrations of 6.3 nM, 10.1 nM, and 38.9 nM and a rapid delayed rectifier potassium current *I*_Kr_ channel block of 75%, 80%, and 90% respectively.

To solve the governing equations 1–6 we adopt the finite element software package Abaqus Unified FEA (part of 3DExperience Simulia software suite, Dassault Systemes, Providence, RI, USA) (1). We exploit the structural similarities of the electrophysiological problem with a heat transfer problem with a non-linear heat source. We discretize the transmembrane potential as a nodal degree of freedom and the ionic currents and gating variables as internal variables (12). Motivated by the small time step size to resolve the fast dynamics during the initial phase of the action potential, we adopt an explicit time integration scheme.

### Cardiac mechanics

To model the mechanical behavior of cardiac tissue, we solve the equilibrium equations derived from Newton’s laws of motion and the conservation of mass. Solving for a static state of equilibrium, these equations translate to

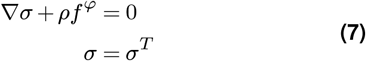

where *σ* is the Cauchy stress tensor, *ρ* is the material density and *f^φ^* is the body force per unit mass. Additional boundary conditions truncate the computational domain. To solve the resulting system of equations, we prescribe constitutive relations between the Cauchy stress *σ* and the tissue deformation and electrophysiology. We first assume the tissue stress state consists of individual passive and active contributions,

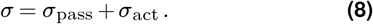

We characterize the kinematics of finite deformation using the deformation gradient

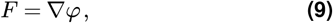

where *ϕ* denotes the deformation field that maps particles *X* in the undeformed material configuration 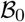 to particles *x* = *φ*(*X,t*) in the deformed material configuration 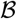. We perform a multiplicative decomposition of the deformation gradient into its volumetric *F*_vol_ and isochoric 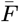 contribution

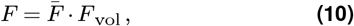

where *F*_vol_ = *J^1/3^I* and the Jacobian *J* = det(*F*). It follows that 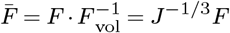. We deduce measures of tissue stretch using the right and left Cauchy-Green tensors, defined as *C* = *F^T^F* and *B* = *FF^T^*. Their isochoric counterparts are defined by 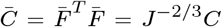 and 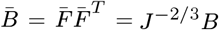. To describe a constitutive stress-stretch relationship that is invariant under superposed rigid body deformations, we define the following stretch invariants

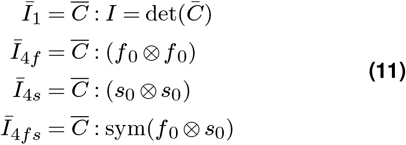

Here *f*_0_ and *s*_0_ describe unit orientation vectors along the considered local material point’s myofiber and sheet direction in the undeformed configuration, respectively. We describe the passive hyperelastic behavior of myocardial tissue using the Holzapfel-Ogden and Arruda-Boyce strain energy function (15; 13; 2). We decompose the strain energy into an isochoric contribution 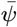 and a volumetric contribution *ψ*_vol_, which reads

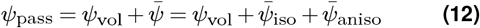

and further decompose the isochoric strain energy into an isotropic and anisotropic contribution. These strain energy contributions read

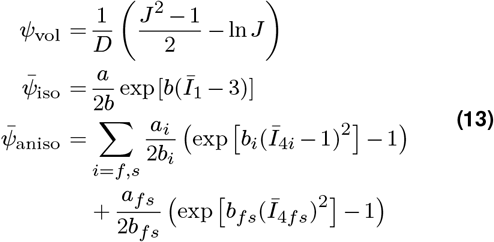

Here 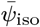 corresponds to the strain energy contributions of the isotropic ground matrix material, whilst 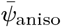 consolidates the anisotropic strain energy contributions of the cardiomyocytes and the families of collagen fibers embedded within the tissue. By deducing the second Piola-Kirchoff stress tensor *S*_pass_ from the strain energy function, we compute the passive Cauchy stress tensor using push-forward operations *σ*_pass_ = *J*^-1^*FS*_pass_F^*T*^,

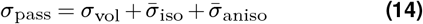

We deduce the following volumetric, isotropic and anisotropic passive Cauchy stress contributions,

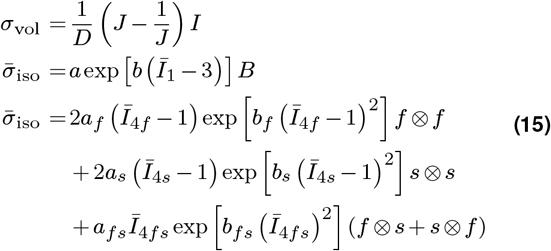

where *f* = *Ff*_0_ and *s* = *Fs*_0_ respectively denote the myofiber and sheet directions in the deformed configuration.

We describe the active stress contribution using a time-varying elastance model (51) which depends on the regional action potential and sarcomere stretch state λ_*f*_ (Frank-Starling effect):

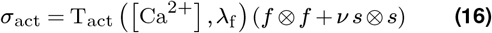

where *ν* describes the active stress interaction between adjacent muscle fibers along the sheet direction *s* (30). The depolarization of the cardiac tissue drives the onset of the active stress generation.

We solve the governing equations 7–16 within the finite element software package Abaqus (1). We set up a Fortran-based user-defined material subroutine describing the Cauchy stress with respect to the deformation invariants, membrane potential (temperature; electrophysiology - heat transfer analogy, ex supra) and time.

### Finite element implementation

The basis for our simulation is the Living Human Heart Model, an anatomically accurate four-chamber model of the healthy human heart (4; 32). The underlying anatomic geometry is based on magnetic resonance imaging of a healthy, 30-year old, 50th percentile U.S. male (54). Images were reconstructed from 0.75mm thick slices using a medium soft-tissue kernel with retrospective electrocardiogram gating. Data acquisition and reconstruction were performed during 70% diastole. The resulting anatomically accurate model includes all four chambers, and the major vessels including the aorta, the pulmonary arteries and the superior vena cava. We prescribe the complex myocardial and atrial architecture of myofiber *f*_0_ and sheet *s*_0_ orientations using rule-based algorithms based on observations from histology and DT-MRI (24; 5; 30).

In this study, we neglect mechano-electrical feedback (40) and successively solve the electrical and mechanical problem. The balance between accuracy and computational cost with respect to element size and critical time step for the defined electrophysiological and mechanical problem leads to two different sets of spatiotemporal discretizations (34; 3). Consequently, we use two different meshes; one to solve the electrophysiological problem in both ventricles specifically and one to subsequently couple the electrophysiological results to the full heart model’s mechanical behavior. For each case, we simulate five seconds without any drug administration followed by an additional five seconds of drug exposure to study the effect of dofetilide on the mechanical behavior and pump efficiency of the heart.

### Electrophysiological drug response

#### Ventricular tissue model

Given our focus on drug-induced ventricular arrhythmogenesis and the fact that the atria are electrically isolated from the ventricles, we concentrate on electrophysiological drug effects in the ventricles. Motivated by the relationship between element size and critical time step size in explicit methods, we converted the ventricular geometry into a regular discretization of cube elements with a constant edge length of 0.3mm across the entire domain. This results in a discretization with 6,878,459 regular linear hexagonal finite elements, 7,519,918 nodes, and 268,259,901 internal variables. For the flux term, we include tissue anisotropy using the fiber definitions *f*_0_ and assign longitudinal and transverse conductivities *D*^||^ = 0.090 mm^2^/ms and *D*^⊥^ = 0.012 mm^2^/ms (26). For the source term, we employ a body flux subroutine to incorporate the ionic currents *I*_ion_ in the solid element formulation (1). To account for and assign regional variations in cell type, we ran a series of Laplace problems using the finite element mesh with different essential boundary conditions (33). From the solutions, we defined the different cell types across the wall, 20% of endocardial cells, 30% of mid wall cells, and 50% of epicardial cells. This arrangement ensures positive T-waves in the healthy baseline electrocardiogram (28).

#### Purkinje network model

The inclusion of the Purkinje network is critical to model correct excitation patterns (20). We create the network as a fractal tree that grows on the endocardial surface (39). This results in a discretization with 39,772 linear cable elements, 39,842 nodes, and 795,440 internal variables. For these Purkinje fiber elements, we developed a linear user element with a discrete version of equation 1. We only connect the Purkinje network to the ventricular tissue at the terminals of the fractal tree (35). For these connections, we use 3,545 resistor elements with a resistance of 1.78*Ω*m, i.e., *χ* = 140mm^-1^ and *C*_m_ = 0.01 *μ*F/mm^2^ (26), between each endpoint of the network and the closest node of the ventricular mesh (6). This allows us to adopt distinct cellular models with different resting potentials for ventricular cells and Purkinje cells. Including resistor elements ensures a bi-directional conduction between Purkinje network and surrounding tissue. For the flux term, we set a conductivity of *D* = 3.0 mm^2^/ms.

#### Electrocardiogram Computation

To calculate pseudo electrocardiograms, at every point 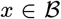 across the heart we project the heart vector ∇*ϕ* onto the direction vector ∇(1/||*r*||) and integrate this projection across the entire cardiac domain 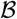 (19; 20),

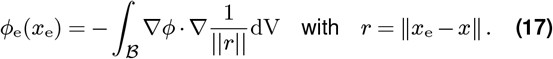

The vector *r* points from current point *x* to the electrode position *x_e_*. To mimic one of the pre-cordial leads in the clinical electrocardiogram, we place the electrode 2cm away from the left ventricular wall. This pre-cordial lead is commonly used to study T waves and QT intervals (14), which are critical to assess the risk of drug toxicity (37).

### Mechanical drug response

#### Electromechanical coupling

For the mechanical problem, a coarser spatial discretization suffices to compute accurate responses (4). Therefore, we discretized the ventricles using 192,040 linear tetrahedral elements with a mean edge size of 2.5 mm and 44,182 nodes. Consequently, the electromechanical coupling requires the interpolation of a three-dimensional 7,519,918 nodal temperature field to a three-dimensional 44,182 nodal temperature field. This was accomplished using Abaqus’s temperature field interpolation functionality between dissimilar meshes in subsequent analyses (1). The full heart mesh, including atria and proximal vasculature parts, comprises 76,282 nodes and 290,723 elements and local fiber- and sheet-orientation assignments. This discretization introduces 228,846 degrees of freedom for the vector-valued deformation. We decribe the atrial action potential, which is not explictly simulated in the electrophysiological ventricular drug-exposure response simulator, using a physiological amplitude step function (18). We report quantitative myofiber stretches across the left and right ventricular wall according to the temporal mean value (and the 95% confidence interval).

#### Coupling to cardiovascular circulation

In order to provide realistic loading conditions and hemodynamic boundary conditions for the atria and ventricles in the heart model, a closed-loop lumped parameter model was set up in Abaqus (4). This lumped parameter model comprises the surface-based fluid cavity representation of the four chambers and additional unit cube fluid cavities representing the arterial and venous systemic and pulmonary circulation respectively. We model the mitral/tricupus valve, the aortic/pulmonary valve, and the systemic/pulmonary resistance flow resistances between these chambers using fluid exchange resistors. We model chamber-specific structural compliances of the additional arterial, venous, and pulmonary chambers using capacitors on one free wall of the unit cube fluid cavities. Since we deduce the geometry of the heart at 70% diastole with the heart already hemodynamically loaded, we estimate the in vivo stress state at the beginning of the simulation using an inverse prestressing method (11; 29).

#### Pressure-volume loops and cardiac output

The pressure and volume in the left and right ventricle is computed using the hemodynamic fluid-cavity definition of both chambers in Abaqus. From these measurements, the pressure-volume loops in both ventricles are extracted. We compute the average stroke volume using the last three simulated dynamically changing pressure volume loops. The average case-specific heart rate is computed based on the average time difference between the last three strokes. Similarly, the time difference between the maximum left and right ventricular contraction is extracted from the last three ventricle-specific contraction sequences. The instantaneous left and right ventricular cardiac output is computed based on the outflow from the left and right ventricular fluid cavity respectively. A 2-second rolling average of this instantaneous outflow expressed as average outflow per minute provides a more descriptive insight on how the cardiac output changes with respect to different administered dofetilide drug concentrations.

## Results

### Electrophysiological drug effects

Figure 2 and the Supplementary Video show the different activation patterns for the baseline case, dofetilide 3x, dofetilide 4.8x and dofetilide 18.5x. These cases correspond to zero, 75, 80 and 90% block of the *I*_kr_ ion channel current respectively. For the baseline case, where no drug is applied, we observe a regular activation sequence that repeats itself ten times in the electrocardiogram. The QRS complex, which represents the fast depolarization driven by the Purkinje network, is preceded by a P wave, which highlights the atrial activation. By blocking the *I*_kr_ current 75%, induced by administration of 3x dofetilide five seconds after drug-free pacing, we see a disruption in the periodic rhythm of the ventricles, leading to arrhythmogenesis that shares features of torsades de pointes. The first electrophysioligcal depolarization wave after drug administration is still driven by the Purkinje network, as shown in the first snapshot, followed by a delay in repolarization, which leads to a secondary activation caused by early afterdepolarizations in a group of midwall cells. The case of 80% block of *I*_kr_ induced by 4.8x dofetilide also shows drug-induced arrhythmogenicity, which is qualitatively similar to the 75% block case. However, the differences in both activation patterns and electrocardiogram recordings highlight the chaotic nature of the arrhythmia, where only a small perturbation in *I*_kr_ block leads to a significantly different temporal evolution of the transmembrane potential. At 3x dofetilide, the left and right ventricle first get activated from base to apex and subsequently from right to left ventricle. At 4.8x dofetilide administration, the depolarization wave evolves towards a left to right ventricular activation sequence. The final case of 90% block of *I*_kr_ caused by 18.5x dofetilide shows an arrhythmia that is closer to ventricular fibrillation, as there are multiple spiral waves driving contractile tissue activation. This chaotic behavior is reflected in the electrocardiogram, where the QRS complexes during the arrhythmia are less defined, with a lower amplitude.

**Figure 1.**
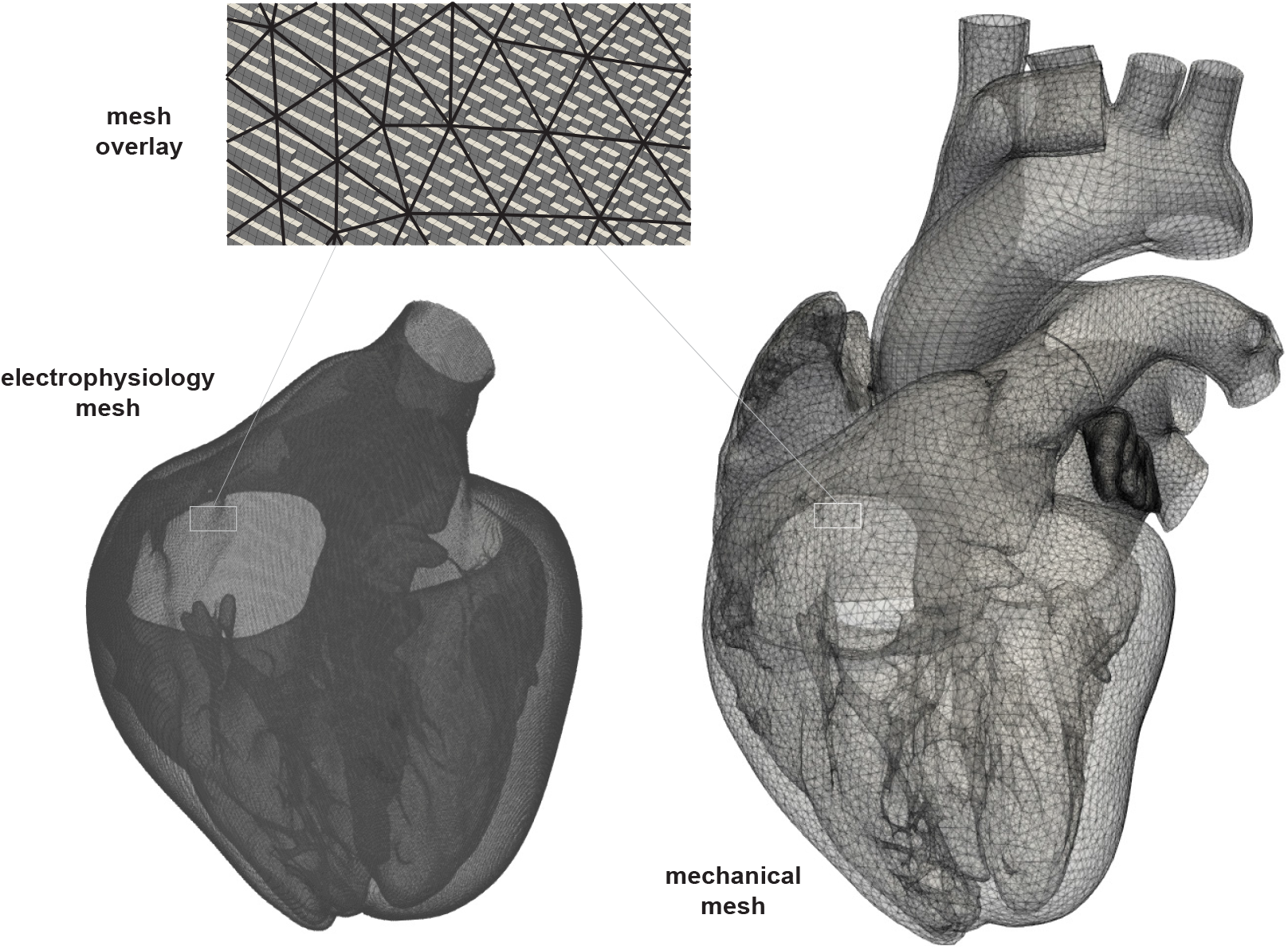
Spatial discretization to compute the electrophysiological and mechanical solution. A mismatch in required spatiotemporal discretization to solve the electrophysiological and mechanical problem leads to two different mesh sizes. To quantify the effects of the drug dofetilide on the activation sequence of the heart, we discretized the ventricles using 39,772 linear cable elements describing the Purkinje fibers and 6,878,459 regular linear hexagonal elements describing the myocardial tissue. Concomitantly, we computed the biomechanical behavior of the ventricles using a mesh consisting of 192,040 tetrahedral elements. We meshed the atria and proximal vasculature using 98,683 additional tetrahedral elements.

**Figure 2.**
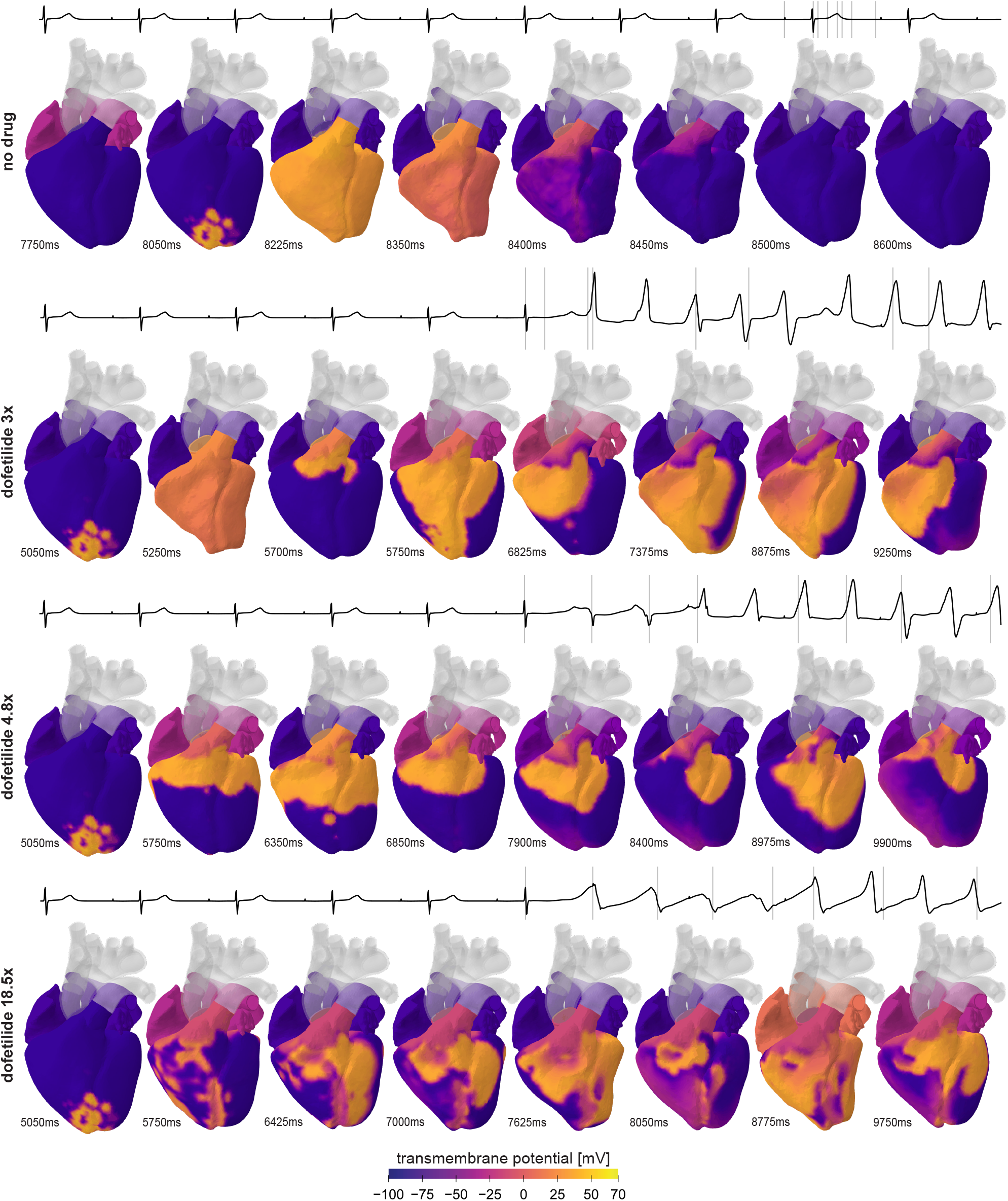
Time evolution of the transmembrane potential for different concentrations of dofetilide. Snapshots are taken at different timepoints for different cases of drug-induced *I*_Kr_ channel block, showcasing the contractile and relaxing deformation in correspondence to the color-plotted electrical activation pattern in the heart. At the top of each row, the computed electrocardiogram signal is shown in black, where the grey vertical lines depict the showcased snapshots for each specific case.

### Mechanical drug effects

Figure 3 and the Supplementary Video highlight the effect that different drug concentrations have on the time sequence of regional myocyte activation over time. The shown snapshots correspond to the time points from Figure 2 with a delay of 50ms (to showcase the locally induced myocardial contraction following a depolarization wave). Figure 4 showacases the left and right ventricular myofiber stretch evolution over time during the five seconds after drug administration. For the baseline no-drug case, the orchestrated depolarization wave of both the left and right ventricle from apex to base causes the ventricles to contract collectively, pushing the blood volume out to the systemic and pulmonary circulation in one cooperative squeeze. More specifically, the myofiber stretches during maximum contraction measure 0.768 (95% CI: 0.651 - 0.898) and 0.711 (95% CI: 0.620 - 0.950) for the left and right ventricle respectively. Moreover, the myocardium is fully relaxed during the atrial contraction, allowing an optimal additional filling of the ventricles during the atrial kick. The myofiber stretches at full relaxation amount to 1.072 (95% CI: 0.981 - 1.165) and 1.058 (95% CI: 0.926 - 1.201) for the left and right ventricle respectively. The myofiber contraction and relaxation remain in complete sync with an average absolute time difference of 25ms between maximum left and right ventricular contraction. For the left ventricle, we compute minimum and maximum myofiber stretches of 0.651 and 1.179 respectively. The 3x dofetilide-induced arrhythmogenicity leads to dissynchronous myocardial contraction and relaxation patterns within the ventricles. Consequently, the myocardial tissue is in active contraction and passive tension at the same time, as can be seen from the wider shaded regions of myofiber stretch variability in Figure 4. In more detail, for dofetilide 3x we compute left and right ventricular myofiber stretches of 0.778 (95% CI 0.661 - 0.918) and 0.723 (95% CI 0.626 - 0.959) at maximum contraction and myofiber stretches of 1.022 (95% CI 0.883 - 1.128) and 0.972 (95% CI 0.804 - 1.100) at maximum relaxation. The drug-induced torsadogenic activation sequence leads to a general right-left ventricular contraction dyssynchrony, during which the right ventricle contracts on average 117ms prior to the left ventricle. Administration of 3x dofetilide leads to minimum and maximum left ventricular myofiber stretches of 0.657 and 1.289. The mechanical effects of 4.8x dofetilide administration are similar to 3x dofetilide, however an important difference between both cases can be found in dyssynchrony. In contrast to 3x dofetilide, upon 4.8x dofetilide administration both left and right ventricular contraction remain synchronized. We compute an average 25ms time difference between left and right ventricular peak contraction, which agrees with the no-drug baseline case. Similar to 3x dofetilide, the left and right ventricular myofiber stretches after 4.8x dofetilide administration amount to 0.777 (95% CI 0.657 - 0.917) and 0.731 (95% CI 0.622 - 0.989) at maximum contraction, and 1.013 (95% CI 0.861 - 1.153) and 0.994 (95% CI 0.817 - 1.145) at maximum relaxation. 4.8x dofetilide affects the minimum and maximum left ventricular myofiber stretches measuring 0.657 and 1.298 respectively. For both 3x and 4.8x dofetilide administration, the maximum myofiber stretches are approximately 10% higher compared to the baseline no-drug case, and typically occur just prior to overall ventricular contraction. This effect arises from the partial contraction of the myocardial tissue during the interventricular pressure buildup phase, causing the tissue that is not activated yet to stretch beyond the baseline physiological stretch range. At the same time, the minimum left ventricular myofiber stretches at 3x and 4.8x dofetilide administration remain relatively comparable to the no-drug baseline case, showcasing the contractile capacity of the tissue is not heavily affected. Upon 18.5x dofetilide administration, the spatiotemporal stretch patterns in Figure 3 are completely irregular, as can be expected from ventricular fibrillation. Consequently, little synchronicity in ventricular contraction and relaxation remains as can be seen from the large shaded temporal myofiber stretch variability shown in Figure 4. The left and right ventricular myofiber stretches amount to 0.858 (95% CI 0.726 - 1.113) and 0.786 (95% CI 0.627 - 1.137) during maximum contraction and 1.017 (95% CI 0.783 - 1.246) and 0.961 (95% CI 0.690 - 1.185) during maximum tissue relaxation. We compute maximum contractile myofiber stretches of 0.715 and maximum relaxing myofiber stretches of 1.262 upon 18.5x dofetilide administration. It should be noted that periodicity in overall ventricular contraction-relaxation fades at this drug concentration, showcased by the smaller amplitude of the mean temporal myofiber stretch evolution and the minimum left ventricular myofiber stretches remaining relatively constant around 0.750 during the last three seconds. During the 18.5x dofetilide-induced ventricular fibrillation, the right-left ventricular dyssynchrony rises to a 92 ms time difference between left and right ventricular peak contraction. Overall, the myofiber stretch variability amounts to a temporally averaged standard deviation of 0.118, 0.114, 0.133 for dofetilide 3x, 4.8x and 18.5x administration respectively. Compared to the no-drug baseline averaged myofiber stretch variability of 0.066, it can be seen how dofetilide affects an effective synchronized contraction of the whole ventricle, and leads to decreasing cardiac pumping efficiency.

**Figure 3.**
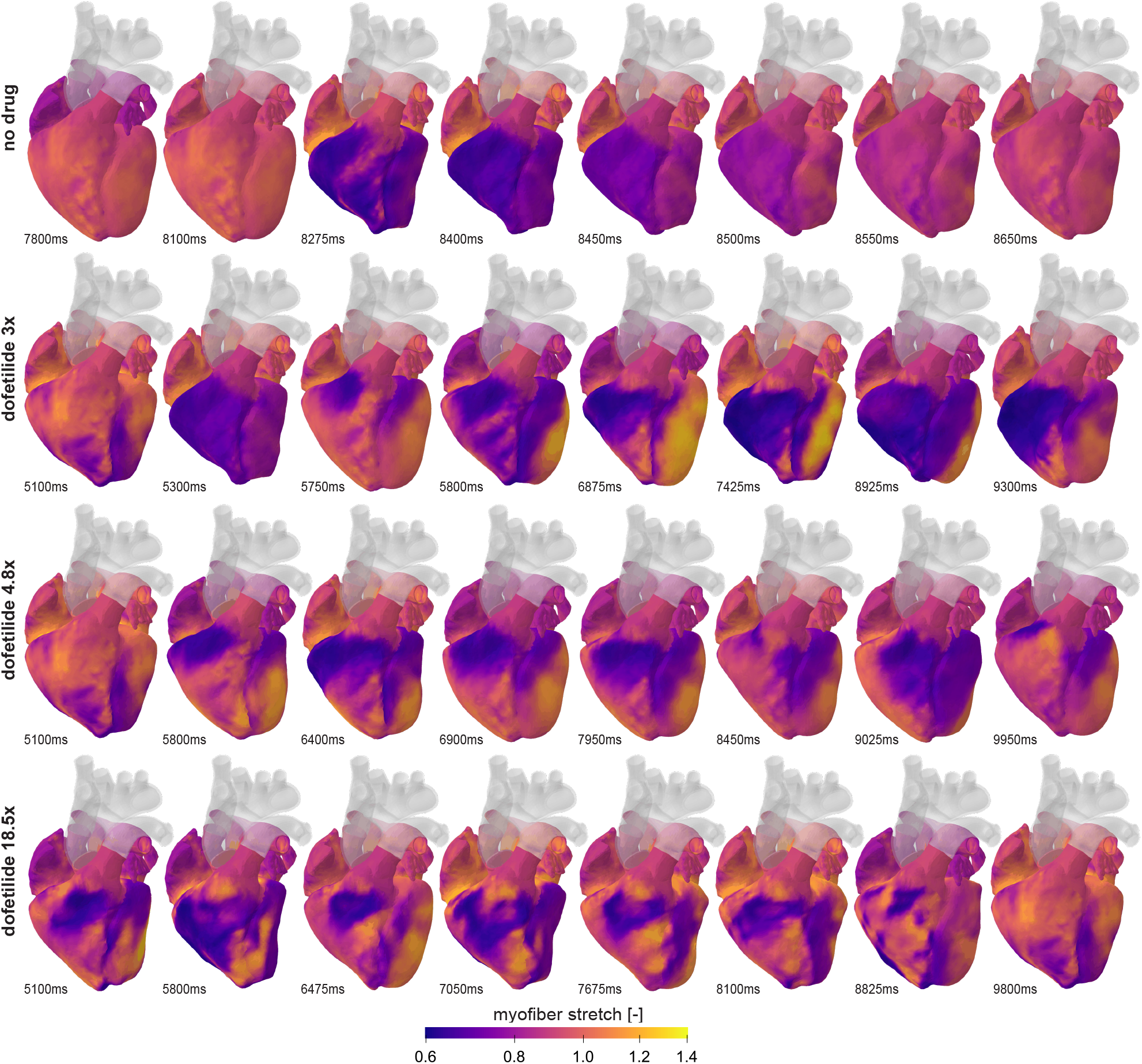
Spatiotemporal evolution of the myofiber stretch for different concentrations of dofetilide. Snapshots are taken at different timepoints for each case, showcasing the effect that blocking of the *I*_Kr_ channel, in correspondence to different administered concentrations of dofetilide, has on the spatiotemporal contraction of the heart.

**Figure 4.**
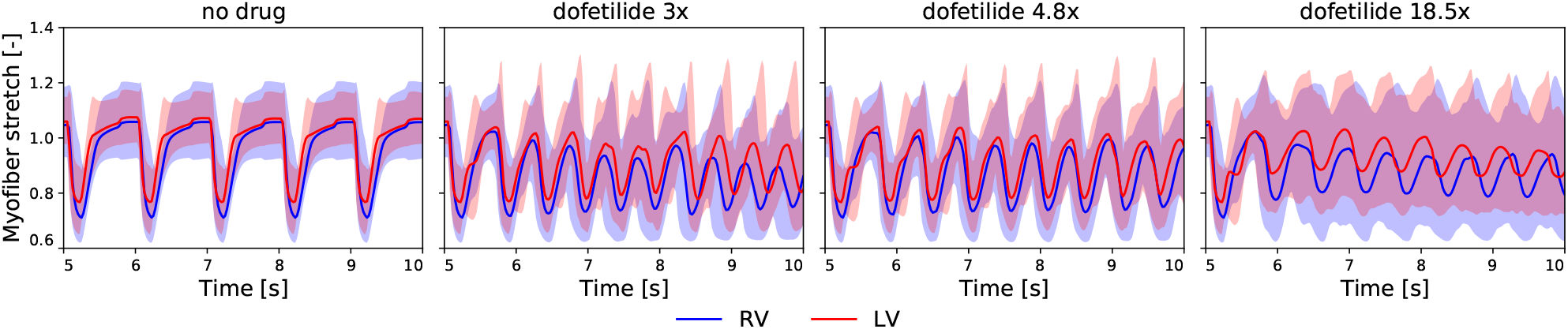
Time evolution of the left and right ventricular myofiber stretch for different concentrations of dofetilide. The temporal myofiber evolution for the left (red) and right ventricle (blue) for each case in correspondence to different administered concentrations of dofetilide. The solid lines showcase the temporal mean value of the myofiber stretch for each respective ventricle whilst the transparent shaded regions represent the ventricle-specific 95% myofiber stretch confidence intervals.

This decreasing cardiac pump efficiency is shown in more detail with respect to the overall cardiovascular circulation in Figure 5. The no-drug baseline pressure-volume loop for the left and right ventricle is shown in the left column. When no drug is administered the stroke volume remains constant at 72 ml. At 3x dofetilide administration, the stroke volume drops to 29 ml and 20 ml for the left and right ventricle respectively. This stroke volume change is mostly caused by a drop in the end-diastolic volume, whilst the end-systolic volume stays approximately the same. The average arrhythmic heart rate after 3x dofetilide administration increases to 123 bpm. At a 80% *I*_Kr_ channel block induced by a 4.8x dofetilide administration, the stroke volume drops from 72 ml to 35 ml for both ventricles. Again, the drop in stroke volume is mostly caused by a smaller end-diastolic volume, whilst the end-systolic volume stays approximately constant. Dofetilide 4.8x causes the average arrhythmic heart rate to increase to 114bpm. At a dofetilide administration of 18.5x, the stroke volume drops to 20 ml for the left ventricle and to 17 ml for the right ventricle. In this case, the drop in stroke volume is caused by both a decrease in the end diastolic volume and an increase in the end-systolic volume. The average arrhythmic heart rate increases to 109 bpm.

**Figure 5.**
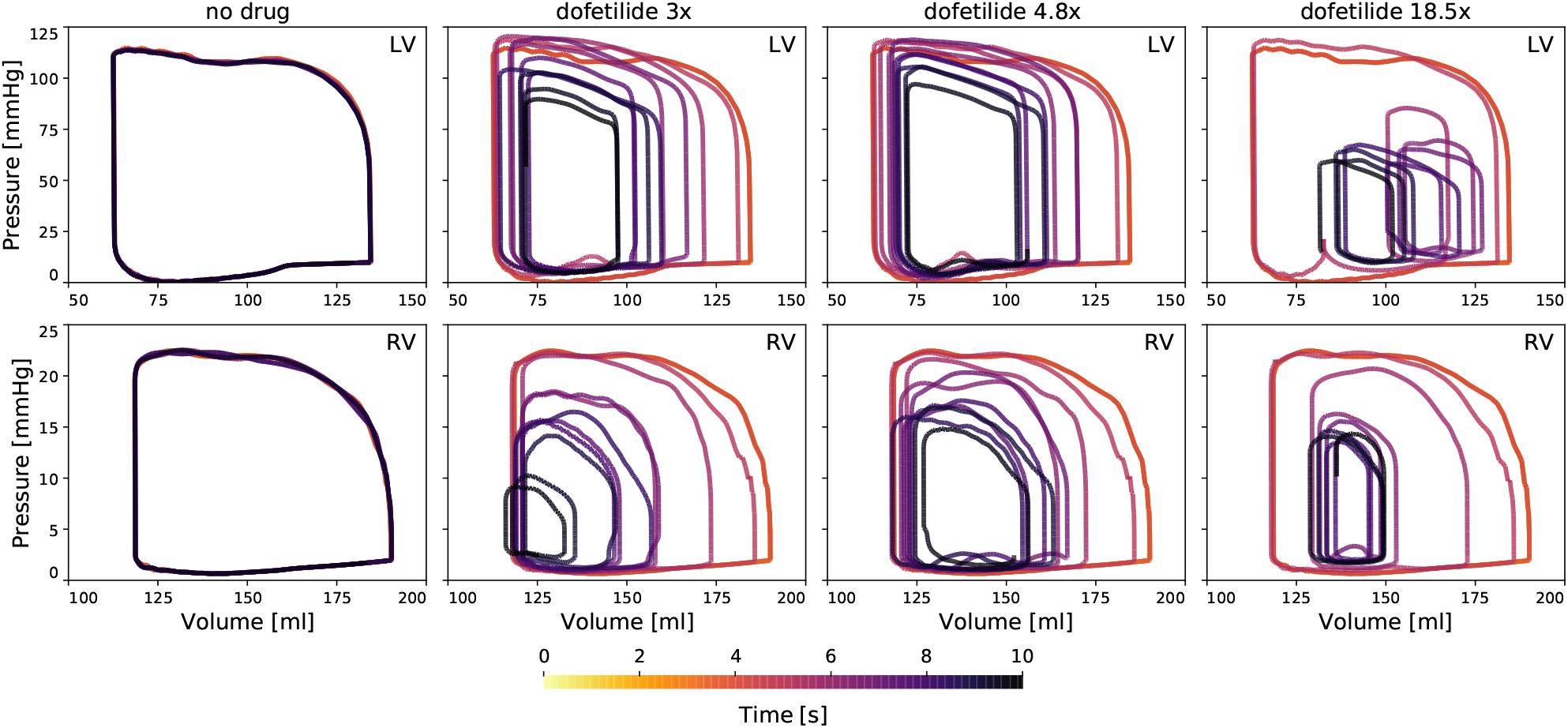
The pharmacological effects of dofetilide on the ventricular pressure-volume loops. Pressure-volume loops showcase the efficiency and frequency of heart contraction for each studied case. For the no-drug case, the pressure-volume loop remains the same. For a 75% *I*_Kr_ channel block (dofetilide 3x), the end-diastolic volume decreases significantly and fluctuates whilst the heart rate increases. For a 80% *I*_Kr_ channel block (dofetilide 4.8x), the end-diastolic volume drops moderately and the heart rate increases. For a 90% *I*_Kr_ channel block (dofetilide 18.5x), the end-diastolic volume drops significantly and the end-systolic volumes increase for both ventricles whilst the heart rate increases.

Figure 6 quantifies the combined effect of drug-induced changing heart rates and stroke volumes on the instantaneous and average cardiac output, denoted by a dotted and solic line respectively, for both the left and right ventricle, highlighted in red and blue respectively. Shown here, 3x dofetilide administration leads to a +5% increase and a −10% decrease in the cardiac output for the left and right ventricle respectively. For 4.8x dofetilide administration, the cardiac output has moderately increased after 5seconds of drug exposure. More specifically, the left and right ventricular cardiac output increased +11% and +3% respectively compared to the baseline cardiac output with no drug exposure. A 18.5x dofetilide administration causes a severe −46% and −64% decrease in left and right ventricular cardiac output respectively.

**Figure 6.**
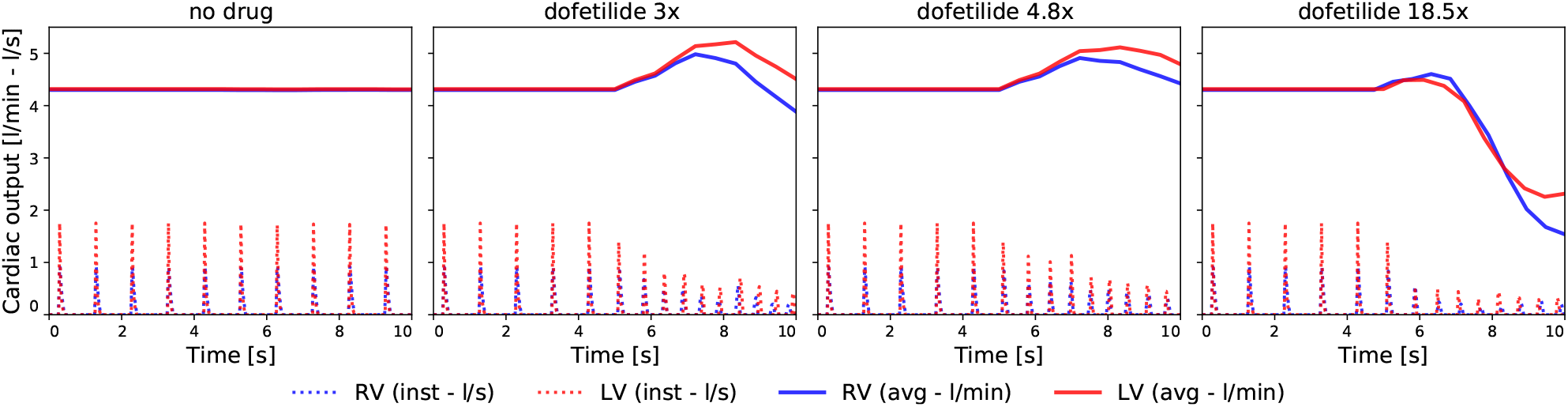
The pharmacological effects of dofetilide on the cardiac output. Cardiac output for the left ventricle (LV - red) and right ventricle (RV - blue) expressed instantaneously (dotted line - l/s) and as a 2-second rolling average (full line - l/min) for the normal case (left column), the mild case (middle column) and the severe case (right column).

## Discussion

Many drugs - not just cardiac drugs - can have serious side effects. One of the most dangerous side effects entails the development of cardiac arrhythmias. More specifically, the development of torsades de pointes - a specific type of polymorphic ventricular tachycardia characterized by a gradual change in amplitude and twisting of the QRS complexes around an isoelectric line on the electrocardiogram (9) - can be especially lethal. Torsades de pointes are often transient but can, in severe cases, lead to ventricular fibrillation causing myocardial damage and even sudden cardiac death. Given its typically short-termed episodic nature, most torsadogenic episodes remain under the radar (17; 50), which leads to limited knowledge on the clinical behavior of the heart during such episodes. When picked up, the clinical evidence of these arrhythmia typically confines itself to electrocardiogram recordings. Pressure-volume loop measurements or flow measurements within a clinical setting are therefore typically unavailable. In this work, we use computational modeling to gather otherwise unattainable insights into the mechanical behavior of the human heart during drug-induced ventricular arrhythmogenicity episodes.

To understand the genesis and development of drug-induced ventricular arrhythmia, cardiac electrophysiology needs to represent both the fast ionic subcellular mechanisms and the slower spatiotemporal cell-tissue-organ scale diffusion process in one and the same framework. To provide accurate physiological outputs and compute potential spiral wave formation, we need a very fine-scaled spatiotemporal discretization of the computational domain (34). Cardiac deformation, on the other hand, is governed by smoother spatial and slower temporal scales. Solving the biomechanical balance equations accurately can therefore be achieved with a much coarser spatio-temporal discretization of the computational domain (3). Given this mismatch in required spatio-temporal discretization and the exponential dependency of computation time on the amount of degrees of freedom to be computed (16), we set up a unidirectional forward electromechanical coupling framework. More specifically, we first computed the electrical propagation of the action potential through the ventricles using a fine-resolution exposure-response simulator (41). Next, the computed spatiotemporal transmembrane potential evolution drives the biomechanical contraction of the cardiac tissue. Given that contractility response of the tissue is critically affected by pre-load and after-load conditions (25), we incorporated an active tension law that depends on the local and temporal sarcomere stretch state λ_*f*_ and coupled the electromechanical heart model to a realistic zero-dimensional surrogate lumped parameter network model of the cardiovascular circulation.

We successfully build an electrophysiological model that inherently captures the regional specificity of the ventricular myocardium and probes the dynamic interplay of its endocardial, midwall, epicardial, and Purkinje cells (42). By extending this model with the dose-dependent effect of dofetilide on the transmembrane ion channel currents, we developed a mechanistic exposure-response simulator that is able to predict the three-dimensional excitation profiles and electrocardiogram recordings shown in Figure 2. The in silico predicted dose-dependent torsadogenic risk of dofetilide agrees favorably with clinical and experimental findings (41; 46). By extending this multiscale framework to a multi-physics framework taking into account the mechanical behavior of the heart and its hemodynamic interaction with the surrounding cardiovascular circulation, we are now able to compute the pharmacological effects that different dosages of dofetilide have on the temporal mechanical behavior of myocardial tissue, as showcased in Figure 3 and 4. Studying the phamarcological effects of different dosages of dofetilide on cardiac pump efficiency involves a complex interplay between regional tissue de- and repolarization, regional tissue contraction and relaxation, and continuously changing hemodynamic loading conditions through the heart’s connection with the surrounding cardiovascular circulation. Therefore, we can only fully appreciate these effects by concomitantly studying the myofiber stretch state in Figure 4, the pressure-volume loops depicted in Figure 5 and the corresponding cardiac output in Figure 6.

For the no-drug baseline case, both ventricles push out the blood in one cooperative synchronized contraction, followed by an extended relaxation phase allowing for atrial blood to refill the ventricle during early diastole and the atrial kick at end diastole. The myofiber stretches cooperatively switch between a contractile and relaxing state, showcased by the low shaded temporal myofiber stretch variability in Figure 4. Naturally, the corresponding pressure-volume loops and cardiac output remain constant.

Upon 3x dofetilide administration, the resulting torsadogenic activation sequence causes an important left-right ventricular contractility dyssynchrony. Immediately after drug administration, the left ventricular contraction starts to trail the right ventricular contraction. Additionally, the drug-induced torsadogenic swirling electrophysiological activation sequence drives the heart rate up to 123 bpm, which causes the ventricles to contract twice before the atria contract. Consequently, the passive atrial-ventricular filling time is significantly shortened, leading to a decreased mean myofiber stretch state during tissue relaxation in Figure 4 and a drop in the left and right ventricular end-diastolic volumes in Figure 5. The corresponding drop in the stroke volume leads to the decreasing instantaneous cardiac output showcased in Figure 6. The increased heart rate also causes an important dyssynchrony between the atrial kick and the ventricular filling phase, further affecting efficient diastolic ventricular filling and decreasing the end diastolic volume. Initially, the atrial kick trails the ventricular contraction, however at specific timepoints within the simulated five-second drug administration timeframe, this dyssynchrony temporarily catches up, as can be appreciated from the fluctuating end diastolic volume evolution in Figure 5. Interestingly, the decreased cardiac output is counterbalanced by the increased heart rate, which partially recovers the expected decrease in cardiac output in Figure 6. For 4.8x dofetilide (80% *I*_Kr_ block), we see a similar combined effect of heart rate and ventricular filling. However, in this case, the torsadogenic activation sequence does not cause a left-right ventricular contraction dyssynchrony. Additionally, a heart rate of 114bpm leads to an atrial kick that leads the ventricular contraction, eventually becoming completely out of phase with the ventricular filling phase at the end of the five second simulated timeframe. This explains the gradual drop in stroke volume in Figure 5. For 3x and 4.8x dofetilide administration, differences in cardiac output in Figure 6 result from dose-dependent interventricular and atrial-ventricular dyssynchronies. Our results showcase a decreased and increased cardiac output for 3x dofetilide and 4.8x dofetilide, highlighting how higher arrhythmogenic drug concentration can impact the cardiac output in a non-intuitive way. Even though the 3x and 4.8x dofetilide induced arrhythmogeneis affects the cardiac output, the depolarization waves swirling around the ventricles still lead to a somewhat temporally structured contraction of the whole ventricle. The resulting active force build up still leads to a decent contraction of the full ventricle, leading to end-systolic volumes that are only slightly larger than for the baseline no-drug case.

For the 18.5x dofetilide case however, the completely chaotic depolarization patterns no longer lead to a synchronous contraction, as can be seen by the large myofiber stretch variability through the whole arrhythmogenic episode in Figure 4. As a result, both the left and right ventricular end-systolic volumes are considerably larger than normal. At the same time, the small re-entrant waves that flicker around the heart also strongly impact the diastolic filling time, leading to decreased end-diastolic volumes. The resulting decrease in stroke volume is so large that the resulting cardiac output in Figure 6 drops significantly. Therefore, the risk for sudden cardiac death at 18.5x dofetilide administration can be expected to be significantly higher than for 3x and 4.8x dofetilide-induced arrhythmia episodes.

Apart from a more mechanistic sudden cardiac death risk stratification, our framework also gives important insights into drug-induced arrhythmogenic overstretching of the tissue. The tissue stretch state is believed to play an important role in pathophysiological growth and remodeling processes (30). Compared to the no-drug baseline case, each drug-induced arrhythmogenic episode showcased increased myofiber stretches in Figure 4. As such, it can be appreciated that our framework provides both acute and chronic mechanistic insights into heart health during and after drug-induced arrhythmogenenesis.

Although our study provides valuable insight into the simultaneous pro-arrhythmic and inotropic liabilities of pharmacological therapies, it has several important limitations that we need to keep in mind when interpreting its results: First, the one-way coupling scheme used in this study does not take into account mechano-electrical feedback (40). It has recently been shown that two-way electromechanical coupling can partially mitigate the action potential duration induced by dofetilide, raising the critical concentration inducing early afterdepolarization onset (25). Second, even though the well-established O’Hara Rudy model used for describing the electrophysiological behavior of the ventricular cardiomyocytes was developed based on a vast amount of human experimental data, a novel update to this model has recently been proposed (49) which reports, amongst others, a re-assessment of the myocardial pro-arrhythmic sensitivity to *I*_Kr_ blockage. Both these limitations might affect the critical drug concentration at which arrhythmia start developing in this study (e.g. 3x dofetilide). Third and final, this study used a unidirectional excitation-contraction model that takes into account myocardial preload and a critical depolarization threshold. Further model development providing a bidirectional coupling between human electrophysiology and active tension generation (21; 40) will allow us to implement more detailed active tension generation models that take into account calcium dynamics, actin-myosin crossbridge cycling transition states and force-frequency responses. Importantly, these coupled models need to remain computationally tractable to be able to compute multiple serial heart beats and potential steady state outcomes. This is a challenging endeavor given the very stiff system of ordinary differential equations for the electrophysiology problem and the amount of state dependent variables in the contraction-excitation coupling, that can require up to 40×2400 CPU hours for simulating one heart cycle, even in a semi-implicit, operator splitted, MPI optimized framework (23). Future work therefore also needs to study the sensitivity of inotropic whole body level results (e.g. end-diastolic and -systolic volumes, ventricular and atrial dyssynchrony, tachycardia) on the biophysical details of the underlying cellular models, and whether or not (potentially machine learning-based) reduced order models can speed up these computations (10).

## Conclusion

This study provides a human-based multiscale and multiphysics mechanistic framework that couples the effect that a drug has on one singular ion channel down at the subcellular level all the way up to a changing cardiovascular circulation at the whole body level. The developed framework provides a granular insight in malignancy of concentration-dependent drug-induced ventricular arrhythmia. Our simulations extend the binary pro-arrhythmic risk classification paradigm for different drug concentrations to an assessment of arrhythmia-severity in light of clinical output metrics as pressure-volume loops and cardiac output. Here, we showed the clinical differences between three drug-induced arrhythmic episodes which results from the fine balance between electrophysiological action potential duration and depolarization times on the one hand and the contractile behavior of the myocardial tissue combined with the contraction of the atria and the connection to the surrounding cardiovascular circulation on the other hand.

## Acknowledgments

This work used the Extreme Science and Engineering Discovery Environment (XSEDE) project TG-MSS170033 supported by the National Science Foundation grant number ACI-1548562, by a Belgian American Education Foundation Postdoctoral Research Fellowship, a Stanford Bio-X IIP Seed Grant, and the National Institutes of Health grant R01HL131975.

## Notes

### Competing Interest Statement

The authors have declared no competing interest.

## References

1. Abaqus Analysis User’s Guide. Dassault Systèmes Simulia Corp., 2020.

2. E.M. Arruda, M.C. Boyce. A three-dimensional model for the large stretch behavior of rubber elastic materials. Journal of the Mechanics and Physics of Solids, 41(2):389–412, 1993.

3. C.M. Augustin, A. Neic, M. Liebmann, A.J. Prassl, S.A. Niederer, G. Haase, and G. Plank. Anatomically accurate high resolution modeling of human whole heart electromechanics: a strongly scalable algebraic multigrid solver method for nonlinear deformation. Journal of Computational Physics, 305:622–646, 2016.

4. B. Baillargeon, N. Rebelo, D.D. Fox, R.L. Taylor, and E. Kuhl. The Living Heart Project: A robust and integrative simulator for human heart function. European Journal of Mechanics. A: Solids, 48:38–47, 2014.

5. J.D. Bayer, C.H. Roney, A. Pashaei, P. Jais, and E.J. Vigmond. Novel radiofrequency ablation strategies for terminating atrial fibrillation in the left atrium: a simulation study Frontiers in Physiology, 7:1–14, 2016.

6. R. Bordas, K. Gillow, Q. Lou, I.R. Efimov, D. Gavaghan, P. Kohl, V. Grau, and B. Rodriguez. Rabbit-specific ventricular model of cardiac electrophysiological function including specialized conduction system. Progress in Biophysics and Molecular Biology, 107(1):90–100, 2011.

7. R. Chabiniok, V. Wang, M. Hadjicharalambous, L. Asner, J. Lee, M. Sermesant, E. Kuhl, A. Young, P. Moireau, M. Nash, D. Chapelle, and D.A. Nordsletten. Multiphysics and multiscale modeling, data-model fusion and integration of organ physiology in the clinic: ventricular cardiac mechanics. Interface Focus, 6:20150083, 2016.

8. W. Crumb, J. Vicente, L. Johannesen, and D. Strauss. An evaluation of 30 clinical drugs against the comprehensive in vitro proarrhythmia assay (CiPA) proposed ion channel panel. Journal of Pharmacological and Toxicological Methods 81:251e262, 2016.

9. F Dessertenne. La tachycardie ventriculaire a deux foyers opposes variables. Archives des Maladies du Coeur et des Vaisseaux, 2(59):263–272, 1966.

10. S. Fresca, A. Manzoni, L. Dedè, and A. Quarteroni. Deep learning-based reduced order models in cardiac electrophysiology. arXiv:2006.03040, 2020.

11. M.W. Gee,, C. Förster, and W.A. Wall. A computational strategy for prestressing patient-specific biomechanical problems under finite deformation. International Journal for Numerical Methods in Biomedical Engineering, 26(1), pp.52–72, 2010.

12. S. Göktepe and E Kuhl. Computational modeling of cardiac electrophysiology: A novel finite element approach. International Journal for Numerical Methods in Engineering, 79(2):156–178, 2009.

13. O. Gültekin, G. Sommer, and G.A. Holzapfel. An orthotropic viscoelastic model for the passive myocardium: continuum basis and numerical treatment Computer Methods in Biomechanics and Biomedical Engineering, 19(15):1647–1664, 2016.

14. J.T.Y. Hii, G. Wyse, A.M. Gillis, H.J. Duff, M.A. Solylo, and L.B. Mitchell. Precordial QT interval dispersion as a marker of Torsade de Pointes. Circulation, 86:1376–1382, 1992.

15. G.A. Holzapfel and R.W. Ogden. Constitutive modelling of passive myocardium: a structurally based framework for material characterization. Philosophical transactions. Series A, Mathematical, physical, and engineering sciences, 367(1902):3445–75, 2009.

16. D.E. Hurtado, and G. Rojas. Non-conforming finite-element formulation for cardiac electrophysiology: an effective approach to reduce the computation time of heart simulations without compromising accuracy. Computational Mechanics, 61(4):485–497, 2018.

17. J. Johnston, S. Pal, and P. Nagele. Perioperative torsade de pointes: a systematic review of published case reports. Anesthesia and analgesia, 117(3):559, 2013.

18. R. Klabunde. Cardiovascular physiology concepts. Lippincott Williams & Wilkins, 2011.

19. M. Kotikanyadanam, S. Göktepe, and E Kuhl. Computational modeling of electrocardiograms: A finite element approach toward cardiac excitation. International Journal for Numerical Methods in Biomedical Engineering, 26(5):524–533, 2010.

20. S. Krishnamoorthi, L.E. Perotti, N.P. Borgstrom, O. Ajijola, A.a Frid, A.V. Ponnaluri, J.N. Weiss, Z. Qu, W.S. Klug, D.B. Ennis, and A. Garfinkel. Simulation methods and validation criteria for modeling cardiac ventricular electrophysiology. PLoS ONE, 9(12):e114494, 2014.

21. S. Land, S.J. Park-Holohan, N.P. Smith, C.G. Dos Remedios, J.C. Kentish, and S.A. Niederer. A model of cardiac contraction based on novel measurements of tension development in human cardiomyocytes. Journal of Molecular and Cellular Cardiology, 106:68–83, 2017.

22. L.C. Lee, J. Sundnes, M. Genet, J.F. Wenk, and S.T. Wall. An integrated electromechanical-growth heart model for simulating cardiac therapies. Biomechanics and Modeling in Mechanobiology, 15(4):791–803, 2016.

23. F. Levrero-Florencio, F. Margara, E. Zacur, A. Bueno-Orovio, Z.J. Wang, A. Santiago, J. Aguado-Sierra, G. Houzeaux, V. Grau, D. Kay, and M. Vázquez. Sensitivity analysis of a strongly-coupled human-based electromechanical cardiac model: Effect of mechanical parameters on physiologically relevant biomarkers. Computer Methods in Applied Mechanics and Engineering, 361:112762, 2020.

24. H. Lombaert, J.M. Peyrat, P. Croisille, S. Rapacchi, L. Fanton, F. Cheriet, P. Clarysse, I. Magnin, H. Delingette, and N. Ayache. Human atlas of the cardiac fiber architecture: study on a healthy population. IEEE Transactions on Medical Imaging, 31(7):1436–1447, 2012.

25. F. Margara, Z.J. Wang, F. Levrero-Florencio, A. Santiago, M. Vázquez, A. Bueno-Orovio, and B. Rodriguez. In-silico human electro-mechanical ventricular modelling and simulation for drug-induced pro-arrhythmia and inotropic risk assessment. Progress in Biophysics and Molecular Biology, 2020.

26. S. Niederer, E. Kerfoot, A.P. Benson, M.O. Bernabeu, O. Bernus, C. Bradley, E.M. Cherry, R. Clayton, F.H. Fenton, A. Garny, E. Heidenreich, S. Land, M. Maleckar, P. Pathmanathan, G. Plank, J.F. Rodríguez, I. Roy, F.B. Sachse, G. Seemann, O. Skavhaug, and N.P. Smith. Verification of cardiac tissue electrophysiology simulators using an N-version benchmark. Philosophical Transactions. Series A, Mathematical, Physical, and Engineering Sciences, 369(1954):4331–51, 2011.

27. T. O’Hara, L. Virág, A. Varró, and Y. Rudy. Simulation of the undiseased human cardiac ventricular action potential: Model formulation and experimental validation. PLoS Computational Biology, 7(5):e1002061, 2011.

28. J. Okada, T. Washio, A. Maehara, S. Momomura, S. Sugiura, and T. Hisada. Transmural and apicobasal gradients in repolarization contribute to T-wave genesis in human surface ECG. American Journal of Physiology - Heart and Circulatory Physiology, 301(1):H200–H208, 2011.

29. M. Peirlinck, M. De Beule, P. Segers, and N. Rebelo. A modular inverse elastostatics approach to resolve the pressure-induced stress state for in vivo imaging based cardiovascular modeling. Journal of the Mechanical Behavior of Biomedical Materials, 85:124–133, 2018.

30. M. Peirlinck, K.L. Sack, P. De Backer, P. Morais, P. Segers, T. Franz, and M. De Beule. Kinematic boundary conditions substantially impact in silico ventricular function. International Journal for Numerical Methods in Biomedical Engineering, 35(1):e3151, 2019.

31. M. Peirlinck, F. Sahli Costabal, K.L. Sack, J.S. Choy, G.S. Kassab, J.M. Guccione, M. De Beule, P. Segers, and E. Kuhl Using machine learning to characterize heart failure across the scales. Biomechanics and Modeling in Mechanobiology, 18:1987—2001, 2019.

32. M. Peirlinck, F. Sahli Costabal, J Yao, J.M. Guccione, S. Tripathy, Y. Wang, D. Ozturk, P. Segars, T.M. Morrison, S. Levine, and E. Kuhl. Precision medicine in human heart modeling: Perspectives, challenges, and opportunities. Biomechanics and Modeling in Mechanobiology (submitted), 2020.

33. L.E. Perotti, S. Krishnamoorthi, N.P. Borgstrom, D.B. Ennis, and W.S Klug. Regional segmentation of ventricular models to achieve repolarization dispersion in cardiac electrophysiology modeling. International Journal for Numerical Methods in Biomedical Engineering, 28:e02718, 2015.

34. S. Pezzuto, J. Hake, and J. Sundness. Space-discretization error analysis and stabilization schemes for conduction velocity in cardiac electrophysiology. International Journal for Numerical Methods in Biomedical Engineering, 32(10):e02762, 2016.

35. A.V.S. Ponnaluri, L.E. Perotti, D.B. Ennis, and W.S Klug. A viscoactive constitutive modeling framework with variational updates for the myocardium. Computer Methods in Applied Mechanics and Engineering, 314:85–101, 2016.

36. J.R. Priest, C. Gawad, K.M. Kahlig, J.K. Yu, T. OHara, P.M. Boyle, S. Rajamani, M.J. Clark, S.T.K. Garcia, S. Ceresnak, J. Harris, S. Boyle, F.E. Dewey, L. Malloy-Walton, K. Dunn, M. Grove, M.V. Perez, N.F. Neff, R. Chen, K. Maeda, A. Dubin, L. Belardinelli, J. West, C. Antolik, D. Macaya, T. Quertermous, N.A. Trayanova, S.R. Quake, and E.A. Ashley. Early somatic mosaicism is a rare cause of long-QT syndrome. Proceedings of the National Academy of Sciences, 113(41):115550–11560, 2016.

37. A. Sadrieh, L. Domanski, J. Pitt-Francis, S. Mann, E.C. Hodkinson, C.A. Ng, M.D. Perry, J.A. Taylor, D. Gavaghan, R.N. Subbiah, J. Vandenberg, and A.P. Hill. Multiscale cardiac modelling reveals the origins of notched T waves in long QT syndrome type 2. Nature Communications, 5:5069, 2014.

38. P.T. Sager. Key clinical considerations for demonstrating the utility of preclinical models to predict clinical drug-induced torsades de pointes British Journal of Pharmacology, 154:1544–1549, 2008.

39. F. Sahli Costabal, D.E. Hurtado, and E Kuhl. Generating Purkinje networks in the human heart. Journal of Biomechanics, 49:2455–2465, 2016.

40. F. Sahli Costabal, F.A. Concha, D.E. Hurtado, and E. Kuhl. The importance of mechano-electrical feedback and inertia in cardiac electromechanics. Computer methods in Applied Mechanics and Engineering, 320:352–368, 2017.

41. F. Sahli Costabal, J. Yao, and E. Kuhl. Predicting the cardiac toxicity of drugs using a novel multiscale exposure–response simulator. Computer Methods in Biomechanics and Biomedical Engineering, 21(3):232–246, 2018.

42. F. Sahli Costabal, J. Yao, and E. Kuhl. Predicting drug-induced arrhythmias by multiscale modeling. International Journal for Numerical Methods in Biomedical Engineering, 34(5):e2964, 2018.

43. F. Sahli Costabal, K. Matsuno, J. Yao, P. Perdikaris, and E. Kuhl. Machine learning in drug development: Characterizing the effect of 30 drugs on the qt interval using gaussian process regression, sensitivity analysis, and uncertainty quantification. Computer Methods in Applied Mechanics and Engineering, 348:313–333, 2019.

44. F. Sahli Costabal, J. Yao, A. Sher, and E. Kuhl. Predicting critical drug concentrations and torsadogenic risk using a multiscale exposure-response simulator. Progress in Biophysics and Molecular Biology, 144:61–76, 2019.

45. F. Sahli Costabal, P. Perdikaris, E. Kuhl, and D.E. Hurtado Multi-fidelity classification using Gaussian processes: accelerating the prediction of large-scale computational models. Computer Methods in Applied Mechanics and Engineering, 357, 112602, 2020.

46. F. Sahli Costabal, K. Seo, E. Ashley, and E. Kuhl. Classifying drugs by their arrhythmogenic risk using machine learning. Biophysical Journal, 118(5):1165–1176, 2020.

47. P. Stewart, O.V. Aslanidi, D. Noble, P.J. Noble, M.R. Boyett, and H. Zhang. Mathematical models of the electrical action potential of Purkinje fibre cells. Philosophical Transactions: Mathematical, Physical, and Engineering Sciences, 367(1896):2225–2255, 2009.

48. K.H. ten Tusscher, D. Noble, P.J. Noble, and A.V Panfilov. A model for human ventricular tissue. American Journal of Physiology: Heart and Circulatory Physiology, 286(4):H1573–89, 2004.

49. J. Tomek, A. Bueno-Orovio, E. Passini, X. Zhou, A. Minchole, O. Britton, C. Bartolucci, S. Severi, A. Shrier, L. Virag, A. Varro, and B. Rodriguez Development, calibration, and validation of a novel human ventricular myocyte model in health, disease, and drug block. eLIFE, 8:e48890, 2019.

50. E. Vandael, B. Vandenberk, J. Vandenberghe, H. Pincé, R. Willems, and V. Foulon. Incidence of Torsade de Pointes in a tertiary hospital population. International Journal of Cardiology, 243:511–515, 2017

51. J.C. Walker, M.B. Ratcliffe, P. Zhang, A.W. Wallace, B. Fata, E.W. Hsu, D. Saloner, and J.M. Guccione. MRI-based finite-element analysis of left ventricular aneurysm. American Journal of Physiology: Heart and Circulatory Physiology, 289(2):H692–H700, 2005.

52. D.W. Wang, I.C. Yang, J.P. Johnson, L. Nie, and P.B. Bennett. Modulation of HERG potassium channels by extracellular magnesium and quinidine. Journal of Cardiovascular Pharmacology, 33(2):181–185, 1999.

53. J. Wong, S. Göktepe, and E. Kuhl. Computational modeling of chemo-electro-mechanical coupling: A novel implicit monolithic finite element approach. International Journal for Numerical Methods in Biomedical Engineering, 29:1104–1133, 2013.

54. Zygote Media Group Inc. Zygote Solid 3D Heart Generations I & II Development Report. Technical Development of 3D Anatomical Systems. 2014.

